# SETD2 safeguards the genome against isochromosome formation

**DOI:** 10.1101/2022.10.25.513694

**Authors:** Frank M. Mason, Emily S. Kounlavong, Anteneh T. Tebeje, Rashmi Dahiya, Tiffany Guess, Logan Vlach, Stephen R. Norris, Courtney A. Lovejoy, Ruhee Dere, Ryoma Ohi, Peter Ly, Cheryl L. Walker, W. Kimryn Rathmell

## Abstract

Factors governing the faithful replication of chromosomes are essential for cellular and genomic integrity. While a variety of mechanisms to manage breaks and promote repair of DNA are widely recognized, epigenetic landmarks that preserve telomere-to-telomere replication fidelity and prevent genome instability are not well-understood. SETD2 is the histone methyltransferase responsible for trimethylation on histone H3 lysine 36 and is newly recognized as a tumor suppressor that acts to maintain genome stability. Importantly, SETD2 is frequently lost in cancers that exhibit extensive intratumoral heterogeneity. Here, we demonstrate that loss of *SETD2* and H3K36me3 promotes chromosome segregation errors and DNA bridging during mitosis, and that these bridges are driven by the formation of dicentric chromosomes. Cytogenetic analyses revealed that these chromosomes were comprised of mirror-imaged isochromosomes and isodicentric chromosomes that contain two active centromeres. These data demonstrate that the SETD2 histone methyltransferase is essential to prevent a palindromic replication intermediate, whose loss precipitates the formation of a mutable chromatin structure known to initiate a cascade of genomic instability in cancer.

## Main

Genome maintenance is one of the highest order functions of a cell. Numerous processes safeguard against errors in replication and segregation of chromosomes during mitosis. Genomic instability and aneuploidy are also hallmarks of cancer and can be initiated by chromosome instability (CIN), defined as a propensity to mis-segregate chromosomes during cell division^1^. Mis-segregated chromosomes can become incorporated into micronuclei, the cytoplasmic DNA-containing structures that are subjected to double-stranded breaks (DSBs) and chromothripsis^2–5^. Amplification or deletion of genomic material or altered chromosomal structures downstream of these events have potential to be deleterious and oncogenic.

One consequence of these chromosomal rearrangements is the generation of chromosomes that contain more than one centromere^6^. Dicentric chromosomes (two centromeres) form chromatin bridges during anaphase, as microtubules from opposite poles attach to both centromeres. Following anaphase, dicentric chromosomes may break causing kataegis and chromothripsis, ultimately promoting subclonal heterogeneity within cell populations and tumors^7–9^. While the consequences of dicentric chromosomes are known, mechanisms that initiate dicentric chromosome formation in most cancer cells are unclear.

SETD2 is a methyltransferase responsible for the tri-methylation of lysine 36 on histone H3 (H3K36me3). SETD2 and this epigenetic mark regulate transcriptional fidelity, RNA splicing, and the DNA damage response^10^. SETD2 is also widely recognized as a tumor suppressor that is frequently mutated or deleted in many tumor types, including clear cell renal cell carcinoma (ccRCC), pediatric acute lymphoblastic leukemia, lung, bladder, and uterine cancer^11^. Intriguingly, recent work in ccRCC demonstrated that tumors with *SETD2* mutations exhibit branched, clonal evolution and more intratumoral heterogeneity than tumors with other driver mutations^12^. Consistent with a role in genome maintenance, *SETD2* loss has been shown to disrupt mitotic spindle organization during mitosis to promote CIN, which may contribute to the tumor suppressive role for this enzyme^13,14^. However, a mechanism linking mitotic errors to the origin of branched evolution in cells or tumors with mutation or deletion of *SETD2*, however, remains unknown.

To quantify the frequency of chromosome mis-segregation following *Setd2* loss, mouse embryonic fibroblasts (MEFs) with wild-type (*Wt*/*Wt*), heterozygous (*Fl*/*Wt*) and homozygous (*Fl*/*Fl*) floxed alleles of *Setd2* were treated with vehicle or 4-OHT to activate Cre-ER and acutely delete *Setd2*. Three days after treatment of 4-OHT, mono-allelic deletion of *Setd2* resulted in decreases in both Setd2 and H3K36me3, whereas bi-allelic loss resulted in complete loss of Setd2 and H3K36me3 (Extended data Figure 1a). H3K36me3 was also lost across chromosome arms and at sub-telomeric and pericentric regions (Extended data Figure 1b). Mono- and bi-allelic loss of *Setd2* increased the prevalence of mitotic cells with lagging, bridging or both types of mis-segregated chromosomes during late anaphase and early telophase (Fig. 1a,b). Lagging chromosomes fail to move poleward, while bridges span across the midzone (Extended data Fig. 1c). To confirm if acute deletion of *SETD2* causes both lagging and bridging chromosomes in human cells, HeLa cells expressing tetracycline-inducible Cas9 (HeLa TetOn-Cas9)^15^, and either non-targeting (NT) or *SETD2*-specific gRNAs, were treated with doxycycline for three days and chromosome mis-segregation quantified. Doxycycline treatment caused loss of SETD2 as well as H3K36me3 along chromosome arms and at sub-telomeric and pericentric regions (Extended data Fig. 1c,d), promoting increases in both lagging and bridging chromosomes (Fig. 1c,d).

**Figure 1:**
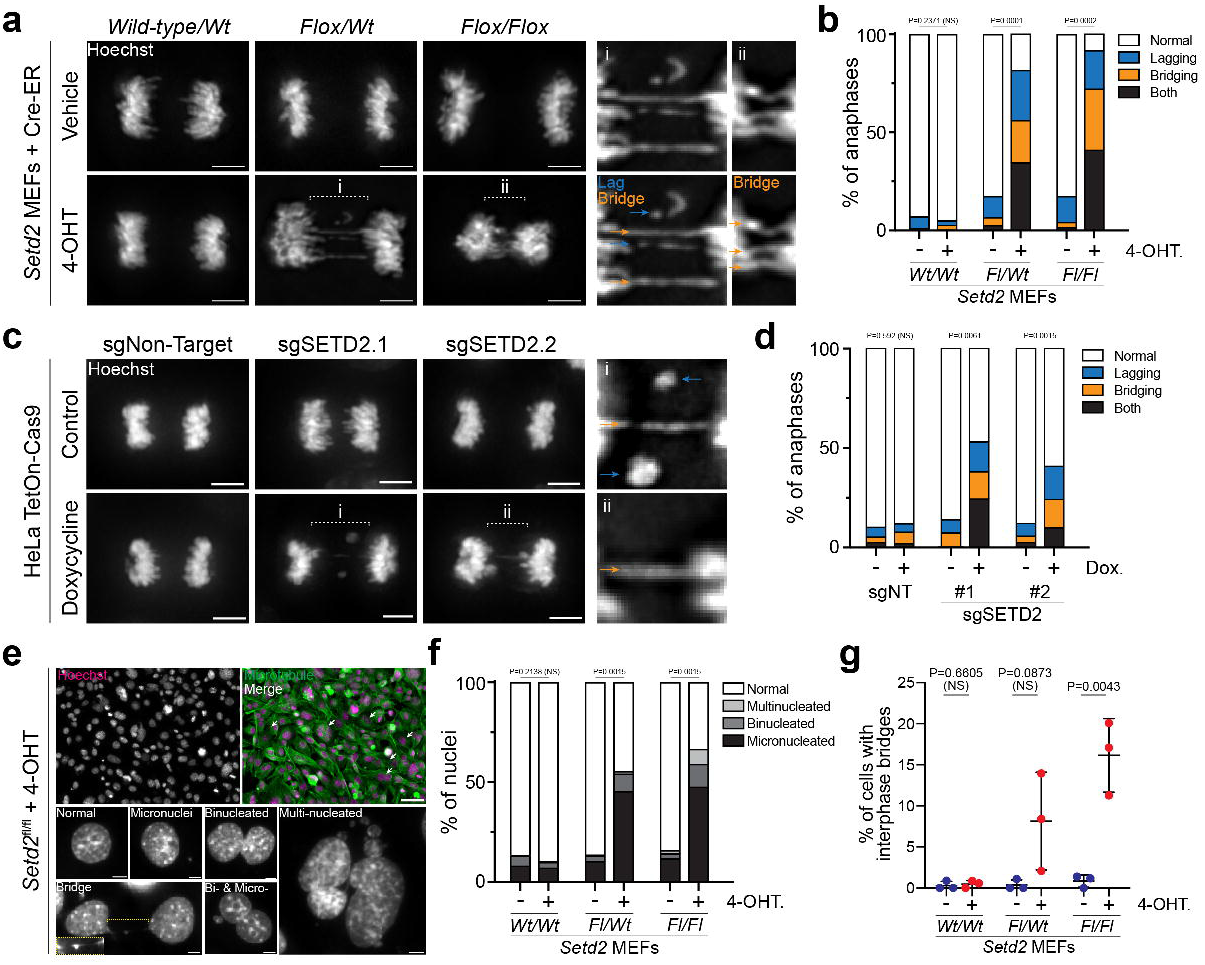
Setd2 loss causes chromatin mis-segregation during mitosis. **a,** Images of wild-type, heterozygous or homozygous deletion of *Setd2* have lagging (blue arrows) and bridging (orange arrows) chromosomes during anaphase. MEFs with no, one or two floxed alleles of *Setd2* (*Wild-type/Wt*, *Flox/Wt*, *Flox/Flox*, respectively) were either treated with Vehicle (EtOH, control in top panels) or 4-OHT (excise floxed alleles of *Setd2*, bottom panels) for three days, fixed and counterstained with Hoechst (grey). Scale bars are 5μm. **b,** Quantification of chromosome segregation errors during anaphase and early telophase in control (vehicle treated) or 4-OHT treated MEFs described in **a**. n=198 *wt/wt* vehicle, n=215 *wt/wt* 4-OHT, n=298 *fl/wt* vehicle, n=227 *fl/wt* 4-OHT, n=258 *fl/fl* Vehicle, n=225 *fl/fl* 4-OHT cells across 2 (*wt/wt*), 4 (*fl/wt*), and 3 (*fl/fl*) biological replicates. P-values derived from unpaired t-test of normal cell values between treatment groups for each genotype. **c,** Images of anaphases in control (untreated) or doxycycline-treated HeLa cells expressing tetracycline-inducible Cas9 (TetOn-Cas9) and single-guide RNA (sgRNA) that are Non-targeting (NT) or specific for *SETD2*. Cells are counterstained with Hoechst (grey). Lagging (blue) and bridging (orange) chromosomes occur in cells lacking *SETD2.* Scale bars are 5μm. **d,** Quantification of chromosome segregation errors during anaphase and early telophase in cells described in **c**. n=183 sgNT control, n=190 sgNT dox., n=198 sgSETD2.1 control, n=240 sgSETD2.1 dox., n=187 sgSETD2.2 control, n=246 sgSETD2.2 dox. cells across 3 biological replicates. P-values derived from unpaired t-test of normal cell values between treatment groups for each sgRNA background. **e,** Image of *Setd2^fl/fl^* MEFs treated with 4-OHT (at left), showing nuclear defects that occur in interphase after three days treatment. Scale bars are 50μm (top images) and 5μm (greyscale images at bottom). **f,** Quantifications of nuclear phenotypes from images described in **e** for *Setd2* MEFs. n=450 *wt/wt* vehicle, n=617 *wt/wt* 4-OHT, n=650 *fl/wt* vehicle, n=509 *fl/wt* 4-OHT, n=736 *fl/fl* Vehicle, n=687 *fl/fl* 4-OHT cells across 3 biological replicates. P-values derived from unpaired t-test of normal cell values between treatment groups each genotype. **g,** Quantification of interphase bridges observed in cell images described in **e**. P-values derived from unpaired t-test of normal cell values between treatment groups each genotype. Error bars are standard deviation (s.d.).

Increased frequency of lagging and bridging chromosomes should lead to an increase in micronuclei and chromatin bridges in interphase^8,9^. Both *Setd2* heterozygous and homozygous-deleted MEFs, as well as HeLa cells acutely deleted of *SETD2*, exhibited increases in binucleation, micronucleation, and multinucleation (Fig. 1e,f and Extended data Fig. 2a,b). Interphase bridges were identified by accumulation of RPA and cGAS, markers of single-stranded DNA and cytoplasmic DNA, respectively (Fig. 1g)^8,16^. Thus, either mono- or bi-allelic loss of *Setd2* is sufficient to promote chromosome segregation defects, which can lead to micronucleation, binucleation, and interphase bridging.

### Loss of *SETD2* causes dicentric chromosome formation

We have previously identified that lagging chromosomes arise from alterations in mitotic spindle morphology following *SETD2* loss, however the defect(s) underlying bridging of chromatin was not clear (Extended data Fig. 1c)^14,17^. Chromatin bridges are caused by structural changes in chromosomes and arise from alterations in chromosome decatenation, cohesion, and/or generation of dicentric chromosomes^9^. To determine whether loss of *SETD2* leads to aberrations in chromosomal structure, metaphase chromosomes from wild-type, mono- and bi-allelic *Setd2* floxed MEFs were isolated three days after 4-OHT treatment, and chromosomal abnormalities were quantified.

Heterozygous or homozygous loss of *Setd2* resulted in an increase in the percentage of cells that have dicentric chromosomes (Fig. 2a,b), as observed by increased DAPI staining, which marks major satellite repeats at centromeres of murine chromosomes^18^. To control for CIN in *Setd2* deleted cells^14^, the number of dicentric chromosome(s) was normalized to chromosome number for each cell, which was confirmed to be increased in *Setd2*-deleted cells (Fig. 2c). 1.1% and 2.0% of chromosomes were dicentric in *Setd2^fl/wt^* and *Setd2^fl/fl^* cells treated with 4-OHT, or 0.43 and 0.81 chromosomes per diploid murine cell, respectively.

**Figure 2:**
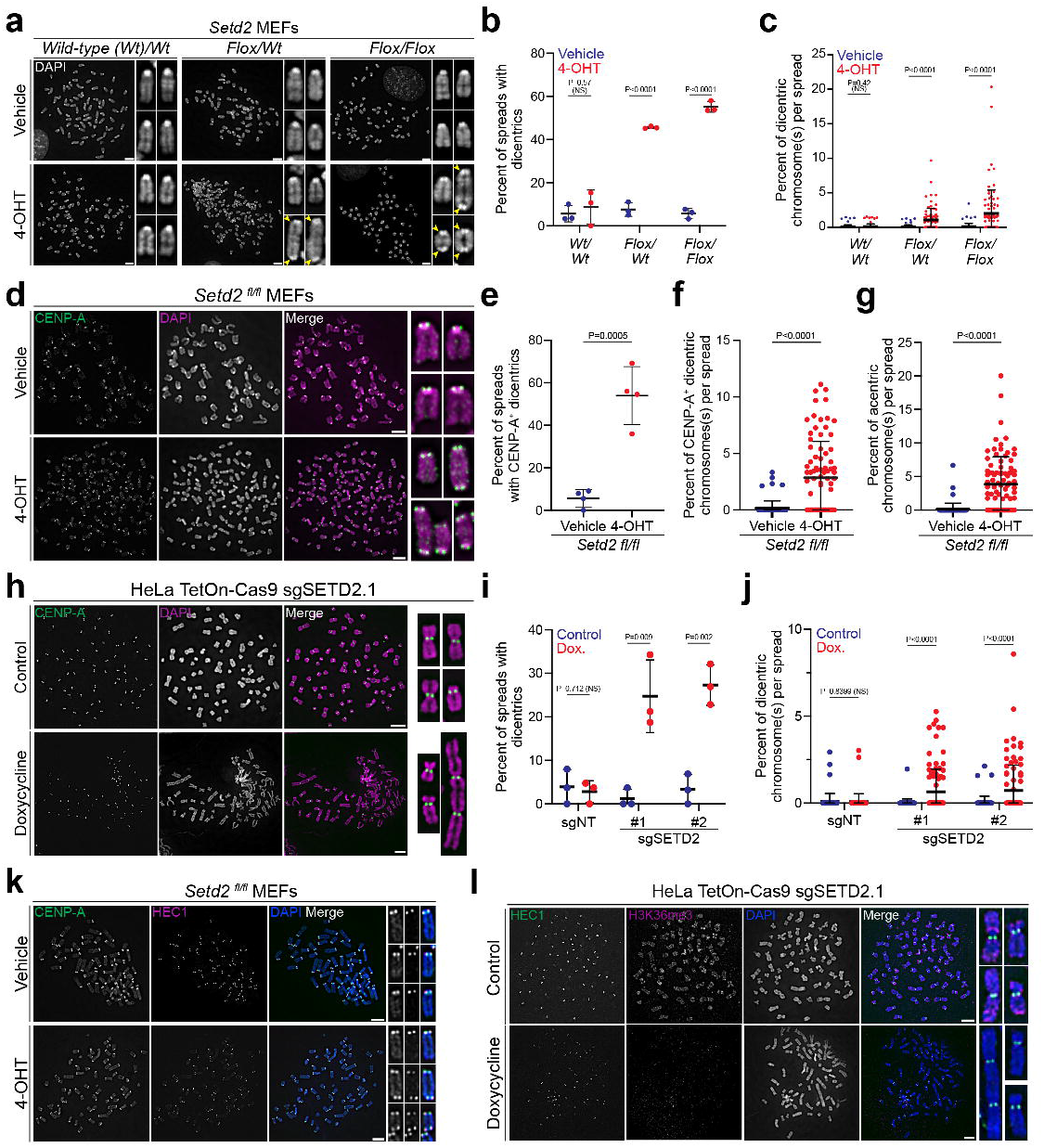
Dicentric chromosomes arise in cells lacking Setd2. **a,** Metaphase spread from MEFs with no, one or two floxed alleles of *Setd2* (*wt/wt*, *fl/wt*, and *fl/fl*, respectively) and treated with Vehicle (control, in top panels) or 4-OHT (bottom panels) for three days. Spreads are counterstained with DAPI (grey). Dicentric chromosomes (yellow arrowheads) are present in cells with homozygous and heterozygous *Setd2* deletion. **b,** Quantification of percentage of metaphase spreads from MEFs described in **a** that have at least one dicentric chromosome from n=3 biological replicates. P-values derived from unpaired t-test of normal cell values between treatment groups each genotype. **c,** Quantification of percentage of dicentric chromosome(s) per metaphase spread image from **b**. Each point represents an individual image. n=88 *wt/wt* vehicle, n=81 *wt/wt* 4-OHT, n=76 *fl/wt* vehicle, n=79 *fl/wt* 4-OHT, n=89 *fl/fl* vehicle, n=85 *fl/fl* 4-OHT cells. P-values derived from unpaired t-test of normal cell values between treatment groups each genotype. **d,** Metaphase spread from *Setd2^fl/fl^* MEFs treated with vehicle (control, top panels) or 4-OHT (*Setd2* knockout, bottom panels). Spreads were immunostained for CENP-A (green), counterstained for DAPI (magenta). Dicentric chromosomes in *Setd2-*knockout cells have two CENP-A foci. **e,** Quantification of percentage of metaphase spreads from *Setd2^fl/fl^* MEFs treated with vehicle (control) or 4-OHT (*Setd2* knockout) that have dicentric chromosomes containing CENP-A from n=4 biological replicates. P-values from unpaired t-test between each condition. **f,** Quantification of percentage of chromosomes that are dicentric and contain two CENP-A foci on metaphase spreads from *Setd2^fl/fl^* MEFs from **e**. n=79 *fl/fl* vehicle, n=85 *fl/fl* 4-OHT cells. P-values from unpaired t-test between each condition. **g,** Quantification of percentage of chromosomes that are acentric from *Setd2^fl/fl^* MEFs treated with Vehicle (n=79 cells) or 4-OHT (n=85 cells). P-values from unpaired t-test between each condition. **h,** Metaphase spread isolated from control (untreated) or doxycycline-treated HeLa TetOn-Cas9 cells expressing sgSETD2.1, immunostained for CENP-A (green) and counterstained for DAPI (magenta). Control cells (top panels) have monocentric chromosomes, while *SETD2*-knockout cells (bottom) have mono- and dicentric chromosomes. **i,** Quantifications of percentage of metaphase spreads that have at least one dicentric chromosome from HeLa TetOn-Cas9 cells expressing sgNT or sgSETD2 and were untreated (control) or treated with doxycycline then stained for CENP-A and DAPI. *SETD2*-knockout caused increases in the frequency of cells that have dicentrics in n=3 biological replicates. P-values derived from unpaired t-test of normal cell values between treatment groups each genotype. **j,** Quantifications of percentage of dicentric chromosomes in each metaphase spread from cells described in **h**. *SETD2-*knockout leads to an increase in the percentage of chromosomes that are dicentric in n=75 sgNT control, n=75 sgNT dox., n=86 sgSETD2.1 control, n=100 sgSETD2.1 dox., n=85 sgSETD2.2 control, n=89 sgSETD2.2 dox. cells. P-values derived from unpaired t-test of normal cell values between treatment groups each genotype. **k,** Metaphase spread from *Setd2^fl/fl^* MEFs treated with vehicle (control, top) or 4-OHT (*Setd2* knockout, bottom), demonstrating that dicentric chromosomes assemble kinetochores (HEC1, magenta) at both centromeres (CENP-A, green). **l,** Metaphase spread from HeLa TetOn-Cas9 cells with sgSETD2, control (top) or treated with doxycycline (*SETD2*-knockout), demonstrating that dicentric chromosomes assemble kinetochores (HEC1, green) at two locations. Scale bars are 5μm. For each plot, error bars are s.d.

To confirm that dicentric chromosomes have two functional centromeres, metaphase spreads from control or *Setd2*-deleted MEFs were immunostained for CENP-A, the histone H3-variant that specifies centromere identity and activity^19^. CENP-A staining verified that 54% of *Setd2*-deleted cells have dicentric chromosomes with CENP-A at two distinct constriction sites on the same chromosome, compared to 6% of controls (Fig. 2d,e). The percentage of chromosomes that are dicentric increased from 0.2% in control samples to 2.8% in *Setd2*-knockout cells (Fig. 2f, 0.08 to 1.12 chromosomes per cell). Additionally, the percentage of acentric chromosomes per metaphase was also increased in *Setd2*-deleted cells (Fig. 2g, 0.2% to 3.9%). We also examined dicentric formation in HeLa TetOn-Cas9 cells with *SETD2* deletion, which also displayed an increase in the percentage of cells with dicentrics (1% to 25% for sgSETD2.1 and 3% to 27% for sgSETD2.2, Fig. 2h,i). The percentage of dicentric chromosomes per spread, as determined by CENP-A staining, also increased in SETD2-knockout cells (0.02% to 0.65% for sgSETD2.1 and 0.06% to 0.72% for sgSETD2.2, Fig. 2j).

For dicentric chromosomes to bridge during mitosis, both centromeres must be active to assemble kinetochores. However, inactivation of either centromere decreases kinetochore assembly, causing the dicentric to segregate like a monocentric chromosome^6^. To determine whether dicentric chromosomes assembled kinetochores at both centromeric locations, metaphase spreads from control or *SETD2*-deleted MEFs and HeLa cells were co-stained for CENP-A and HEC1, the outermost component of the kinetochore that binds microtubules. Indeed, both centromeres of dicentric chromosomes were fully active (Fig. 2k,l), indicating that these dicentrics contribute to the elevated frequency of anaphase bridges induced by SETD2 loss.

### Inter-chromosomal rearrangements, telomere fusion and neocentromere formation are not primary drivers of dicentric chromosomes following *Setd2* loss

Dicentric chromosomes are a product of chromosome fusion or neocentromere formation and are known to occur in human diseases and promote genomic instability and tumor evolution^7,9,20,21^. A common driver of chromosome fusion is telomere dysfunction, and H3K36me3 is present near telomeric or sub-telomeric regions (Extended data Fig. 1b,e)^8^. To test whether telomere attrition occurs following acute *Setd2* loss, quantitative fluorescence *in situ* hybridization (Q-FISH) was performed to determine telomere length in vehicle and 4-OHT treated *Setd2^fl/fl^* MEFs. There were no changes in telomere length associated with loss of *Setd2* (Extended data Fig. 3a-c). Additionally, no telomeric FISH signal was observed between the two centromeres, indicating that telomere fusion was not occurring (Extended data Fig. 3d,e). Together, these results indicate that telomere dysfunction is not the primary driver of dicentric chromosome formation following acute loss of *Setd2*.

We next examined whether dicentric chromosomes arise due to neocentromere formation or telomere-independent chromosome fusion, which may be downstream of DSBs. If chromosome fusion drives dicentric formation, those chromosomes should harbor an inter-chromosomal rearrangement between two non-homologous chromosomes. However, if dicentrics lack inter-chromosomal rearrangements, this may suggest neocentromere formation. To differentiate between these possibilities, multicolor FISH (m-FISH) was performed on metaphase spreads from vehicle or 4-OHT treated *Setd2^fl/fl^* MEFs to determine the genetic composition of the dicentric chromosomes (Fig. 3a,b). m-FISH identified that 15.3% of dicentrics have inter-chromosomal rearrangements, generated from the fusion of either two or more different chromosomes. However, the majority (84.7%) of dicentric chromosomes do not have inter-chromosomal rearrangements and are comprised of the same chromosome (Fig. 3c). Moreover, there is no preference for dicentric formation on specific chromosomes.

**Figure 3:**
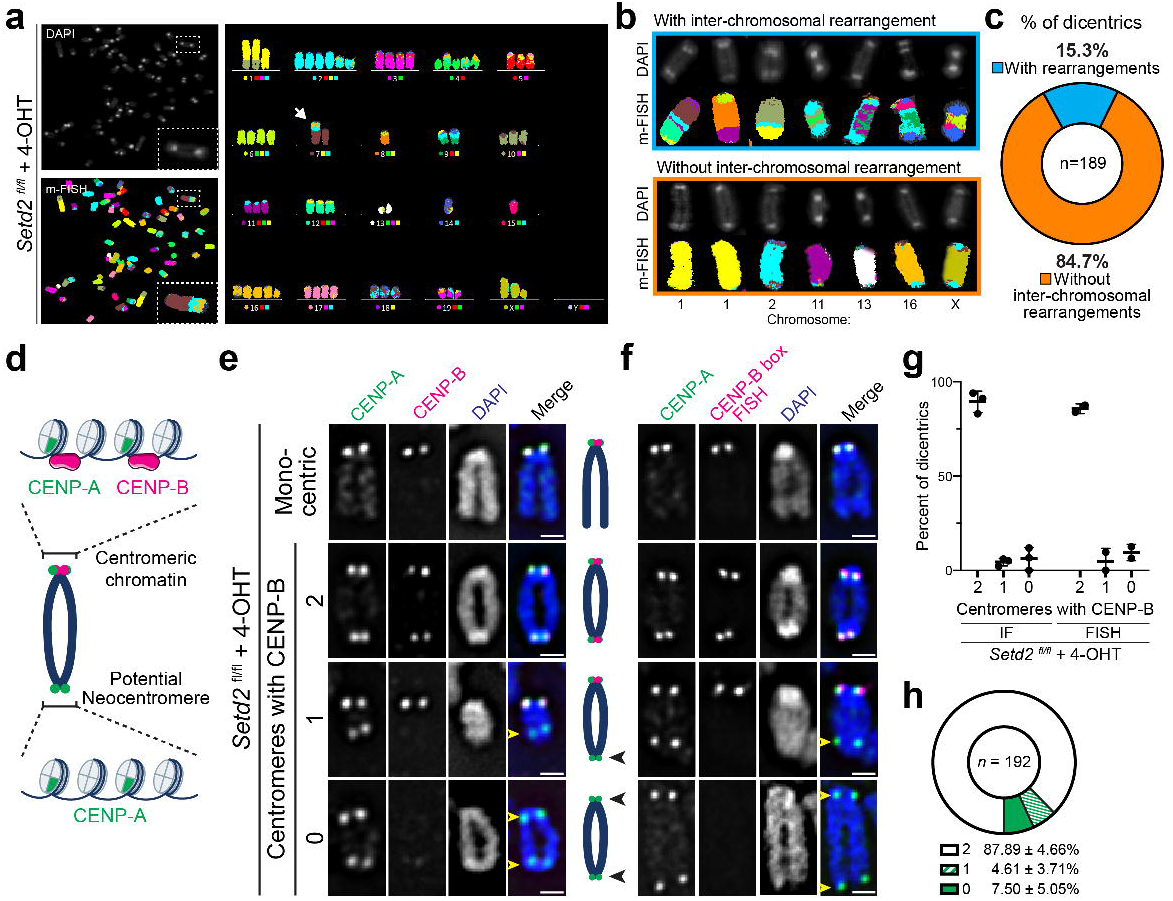
Dicentric chromosomes in Setd2-knockout cells lack inter-chromosomal rearrangements and neocentromeres. **a,** Multicolor fluorescence *in situ* hybridization (m-FISH) image on *Setd2^fl/fl^* MEFs treated with 4-OHT (*Setd2*-knockout). Arrow indicates dicentric chromosome. **b,** Representative images of dicentric chromosomes with inter-chromosomal (top panels) or intra-chromosomal rearrangements (bottom panels). **c,** Quantification of dicentric chromosomes (n=189 across three biological replicates) with or without inter-chromosomal rearrangements. The majority of dicentrics lack inter-chromosomal rearrangements and are comprised of genetic material from the same chromosome. **d,** Schematic of dicentric chromosomes. CENP-B binds CENP-B box motif on centromeric DNA. Neocentromeres, which contain CENP-A nucleosomes, would lack CENP-B. **e,** Representative immunofluorescent images of chromosomes from *Setd2^fl/fl^* MEFs treated with 4-OHT and stained for CENP-A (green), CENP-B (magenta), and DNA (DAPI, blue). CENP-A localizes with CENP-B at centromeres in monocentric cells and in some dicentrics. Chromosomes lacking CENP-B on one or both centromeres (yellow arrowheads) may be suggestive of neocentromeres. Scale bars are 1μm. **f,** Representative immunofluorescent and fluorescence *in situ* hybridization (FISH) images of chromosomes from *Setd2^fl/fl^* MEFs treated with 4-OHT and stained for CENP-A (green), CENP-B box FISH probe (magenta), and DNA (DAPI, blue). CENP-A localizes with CENP-B box DNA at centromeres in monocentric cells and in some dicentric cells. Some centromeres contain CENP-A but lack CENP-B box sequences (yellow arrowheads). Scale bar is 1μm. **g,** Quantifications of percent of dicentrics that have CENP-B co-localizing with CENP-A at 0,1, or 2 sites across n=3 (IF) and n=2 (FISH) biological replicates. Error bars are s.d. **h,** Quantification of CENP-A co-localization with CENP-B from n=192 dicentrics described in **c**. Mean percentage (± s.d.) across 5 biological replicates (IF and FISH).

Dicentrics may occur by establishing neocentromeres at an ectopic, non-centromeric site^22^. To test whether neocentromeres form in *Setd2*-knockout cells, metaphase spreads from vehicle and 4-OHT treated *Setd2^fl/fl^* MEFs were immunostained for CENP-A and CENP-B (Fig. 3d,e), the latter of which binds to CENP-B boxes, a 17 base-pair motif sequence located within native a-satellite-containing centromeres for all chromosomes (except the Y-chromosome)^23^. Using a complementary approach, spreads were immunostained for CENP-A, followed by FISH using a CENP-B box probe (Fig. 3F). If centromeres form at the correct location, CENP-A should co-localize with CENP-B. If neocentromeres form at ectopic sites, CENP-A would be present at sites lacking CENP-B (Fig. 3d). The percentage of centromeres localizing with CENP-B was quantified, using either immunofluorescence to detect CENP-B or FISH against the CENP-B box DNA (Fig. 3g). Using either approach, both centromeres contained CENP-B in 88% of dicentrics, and 12% of dicentrics contained CENP-A at potential non-centromeric sites (Fig. 3h). It should be noted that the *Setd2^fl/fl^* MEFs were derived from a male mouse, and the Y-chromosome lacks a CENP-B box. Thus, we cannot differentiate between Y-chromosome-derived dicentrics and potential neocentromeres. While these observations indicate that *Setd2* deletion may possibly promote some degree of neocentromere formation, the vast majority of dicentric chromosomes are not formed by this mechanism.

### Isodicentric chromosomes predominate in *Setd2*-deleted cells

We examined the nature of the dicentrics in *Setd2*-knockout cells more carefully. One possibility was that the dicentrics were isodicentric, defined as a derivative chromosome comprised of a mirror-imaged duplication that promotes gene amplification and deletion within an individual chromosome^24^. Isodicentrics were suspected because centromeres were exclusively positioned at both chromosome poles in *Setd2*-deleted MEFs, suggesting symmetry (Fig. 4a). Dicentrics in *SETD2*-knockout HeLa cells also appeared symmetrical, often producing abnormally large chromosomes (Fig. 4b). To test whether isodicentric chromosomes occur in *Setd2*-mutant MEFs, metaphase spreads were subjected to Geimsa staining and G-banding analyses (Fig. 4c). This cytogenetic approach identified 75% of dicentrics as bona fide isodicentric chromosomes.

**Figure 4:**
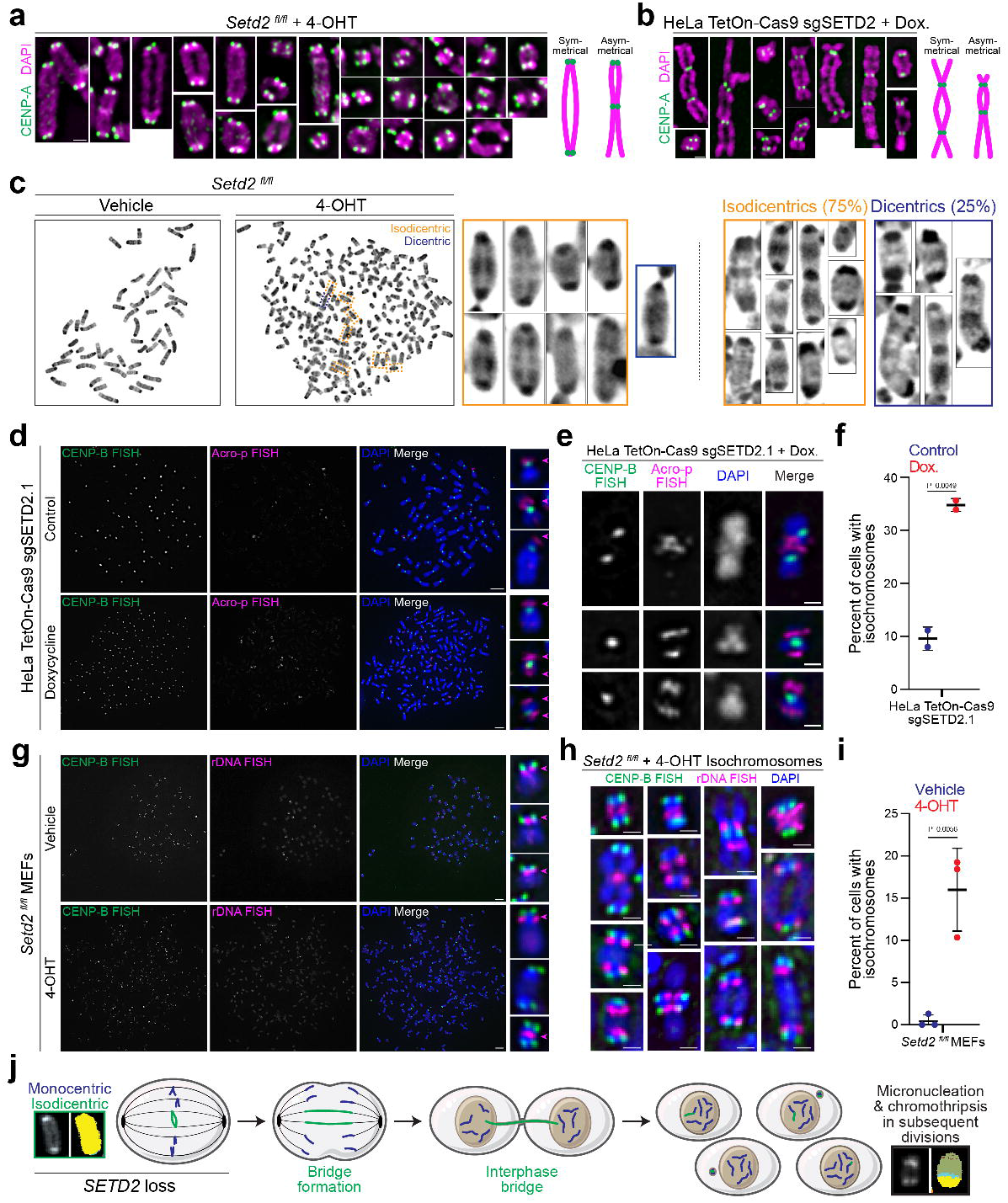
Setd2 knockout promotes formation of isodicentric chromosomes. **a,** Immunofluorescent images of dicentric metaphase chromosomes from *Setd2^fl/fl^* MEFs treated with 4-OHT (*Setd2*-knockout) and stained for CENP-A (green) and DNA (DAPI, magenta). Nearly all dicentric chromosomes are symmetrical. Scale bar is 1μm and all images are set to same scale. **b,** Representative immunofluorescent images of dicentric metaphase chromosomes from HeLa TetOn-Cas9 cells expressing sgSETD2.1 and treated with doxycycline (*Setd2*-knockout). Chromosomes are stained for CENP-A (green) and DNA (DAPI, magenta). Scale bar is 1μm. **c,** Images of Geimsa stained metaphase spreads from *Setd2^fl/fl^* MEFs treated with vehicle (control) or 4-OHT (*Setd2*-knockout). Cytogenetic analyses identified that vast majority of dicentrics are isochromosomes (isodicentric, orange boxes). Dicentric chromosomes (blue boxes) occur but the majority are isodicentric. n=24 dicentrics across 18 spreads. **d,** Images of FISH for CENP-B box and p-arms of acrocentric chromosomes (Acro-p) on metaphase spreads from control or doxycycline-treated HeLa TetOn-Cas9 cells expressing sgSETD2.1. Acro-p signal should be at short, p-arm (top panels), however *SETD2*-knockout cells have signal on both arms. **e,** Representative images of isochromosomes, including isodicentric, from HeLa TetOn-Cas9 sgSETD2.1 cells treated with doxycycline. **f,** Quantification of percentage of cells with isochromosomes in control or doxycycline-treated HeLa TetOn-Cas9 sgSETD2. n=2 biological replicates. **g,** Images of FISH for CENP-B box and rDNA (BAC RP23-225M6) on metaphase spreads from control or 4-OHT-treated *Setd2^fl/fl^* MEFs. rDNA should be present at one location per chromosome, however *Setd2*-knockout leads to isodicentrics (bottom right). **h,** Representative images of isochromosomes, including isodicentrics, from *Setd2*-knockout MEFs. **i,** Quantification of percentage of cells with isochromosomes in control (159 spreads) or 4-OHT-treated (241 spreads) *Setd2^fl/fl^* MEFs across n=3 biological replicates. **j,** Proposed model describing how *SETD2* loss drives genetic heterogeneity. Isodicentric chromosomes initiate gene amplification and deletion, chromosome mis-segregation, and chromothripsis in subsequent divisions to increase clonality. For each plot, error bars are s.d. and P-values derived from unpaired t-test.

To confirm whether isochromosomes occur in *SETD2*-deleted cells, FISH against nucleolar organizer regions was performed to simultaneously mark the p-arms of the human acrocentric chromosomes (13, 14, 15, 21, and 22, Acro-p) or 45S rDNA of mouse chromosomes (11, 12, 15, 16, 18, 19). While control HeLa samples showed stereotypic Acro-p FISH staining, *SETD2*-knockout chromosomes exhibited more cells with isochromosomes, including isodicentrics (Fig. 4d-f). Formation of isochromosomes and isodicentrics were also observed in *Setd2*-knockout MEFs (Fig. 4g-i).

While dicentric chromosomes arise from faulty DSB repair, telomere attrition, or neocentromere formation^20^, mechanisms promoting isodicentrics are less well understood. In yeast, mirror-image dicentric and acentric chromosomes can arise through a DNA replication-dependent, DSB-independent manner called faulty template switching or replication template exchange^25,26^. During this process, stalled replication forks restart utilizing RAD52 instead of the recombinase RAD51 and anneal to a nearby inverted repeat, essentially performing “U-turns” to generate palindromic acentric and dicentric chromosomes. Loss of H3K36me3 decreases RAD51-dependent homologous recombination^27^, so one hypothesis was that cells lacking SETD2 might preferentially utilize RAD52 during DNA repair. To test this, control or *SETD2*-knockout HeLa cells were irradiated (5Gy), and immunostaining for RAD51 and RAD52 revealed that cells lacking SETD2 had decreased RAD51 foci and more RAD52 foci (Extended data Fig. 4a-d). While total protein levels of these factors were not affected (Extended data Fig. 4e), *SETD2*-knockout cells demonstrated increased loading of RAD52 onto chromatin following irradiation (Extended data Fig. 4f). These data suggest that cells lacking H3K36me3 and SETD2 may utilize an alternative homologous recombination pathway that is more prone toward faulty template switching.

## Discussion

SETD2 is the sole source of H3K36 trimethylation and a known tumor suppressor, with roles in mitotic spindle organization, DNA repair and transcriptional fidelity. Yet, it has been unclear how *SETD2* loss causes extensive intratumor heterogeneity and branched evolution. We propose that, in addition to CIN, the formation of isochromosomes following *SETD2* loss of function may provide a mechanism to promote the genomic heterogeneity observed (Fig. 4j). This model is consistent with previous work demonstrating that a single chromatin bridge is sufficient to promote clonal heterogeneity^7,9^, suggesting that this structural intermediate may in fact be the major feature driving genomic instability.

Chromosome bridges are subject to cycles of breakage-fusion-bridge (BFBs), which initiate further gross chromosomal rearrangements and chromothripsis in subsequent cell divisions^9,28^. Genomic signatures of BFBs occur in myriad tumor types, and dicentric chromosomes are sufficient to promote cellular transformation^21,29,30^. Yet, it is unclear what molecular or genetic changes initiate dicentric chromosome formation. Our findings establish that SETD2 prevents this mutable chromosomal structure, and that *SETD2* loss is sufficient to promote isodicentric chromosome formation and bridging, a critical step to initiate BFB, gross chromosomal rearrangements and clonal heterogeneity.

The ability to generate more clones provides tumors a fitness advantage, and accordingly, a range of tumor types with *SETD2* mutations are more resistant to a broad range of therapeutics^31–34^. Notably, ccRCC, where *SETD2* was initially established as a tumor suppressor, is among the most resistant cancers to chemotherapy^35^. *SETD2* mutations commonly occur in hepatosplenic T-cell lymphoma (HSTL), a rare yet aggressive neoplasm that is also chemoresistant^36^. The most common chromosomal abnormality in HSTL is isochromosome 7q [i(7q)], which promotes amplification of genes in 7q and deletion of 7p. Further analyses are needed to assess whether isochromosomes occur in other tumor types lacking SETD2.

These palindromic or mirror-image chromosomes promote copy-number variations, gene amplifications, and gross chromosomal rearrangements observed in cancer and genetic disorders^24,37^. While faulty template switching and RAD52 have been shown to drive “U-turns” in yeast, fork stalling and template switching (FoSTeS) and microhomology-mediated break-induced replication may also promote isochromosomes, leading to copy number alterations and aneuploidy in mammalian systems^38,39^. Lastly, non-allelic homologous recombination (HR) between sister chromatids causes isodicentric chromosome formation associated with Turner syndrome, sex reversal and spermatogenesis defects^40^. While it is unclear which of these mechanisms may occur here, *SETD2* loss may prime cells for palindromic rearrangements because SETD2 and H3K36me3 are required for DNA replication fidelity, DSB repair, and HR.

These data demonstrate that loss of the tumor suppressor *SETD2* promotes the formation of isochromosomes, acentric, dicentric, and isodicentric chromosomes, which form chromatin bridges during mitosis. While these structures have long been appreciated to promote intratumoral heterogeneity, their origin has been elusive. Our data provides an unexpected and important link between an essential histone methyltransferase and genome maintenance.

## Methods

### Cell culture

All cells were cultured at 37°C and 5% CO_2_. Mouse embryonic fibroblast (MEFs) were cultured as described previously^14^. Briefly, cells were cultured in DMEM, phenol red-free (Gibco, Catalog number 21063-029) supplemented with 1mM sodium pyruvate (Sigma, S8636-100mL), GlutaMAX (Gibco, 35050-061), and 10% fetal bovine serum (FBS, GeminiBio, 100-106). HeLa TetOn-Cas9 and HEK293FTs cells were grown in DMEM (Gibco, 11965-092) supplemented with 1% Penicillin/Streptomycin (Gibco, 15150122) and 10% tetracycline-free FBS (GeminiBio, 100-800)^15^. *Setd2^wt/wt^, Setd2^flox/wt^, Setd2^flox/flox^* MEFs expressing ER-Cre were treated with 3μM (Z)-4-hydroxytamoxifen (4-OHT, Millipore Sigma, H7904) or vehicle (Ethanol, 0.1% final volume) for three days, unless stated otherwise, then harvested for experiment.

### Cell culture methods, treatments and cell line generation

Single guides targeting *SETD2* (sgSETD2.1 sense strand GTGCGGATCAGCCAATTGCCG and sgSETD2.2 sense strand GAATGAACTGGGATTCCGACG) were cloned into pLenti-sgRNA (Addgene, 71409) following previous methods^41^. Lentivirus was generated using 293FT cells by co-transfection of guide plasmid with psPAX2 and pMD2.G (Addgene, 12260 and 12259, respectively). Conditioned media was harvested, filtered, and stored at −80. HeLa TetOn-Cas9 cells were transduced with 8mg/mL polybrene (Millipore Sigma, TR-1003) and viral supernatant for guides targeting *SETD2* or non-targeting control sgRNA (Addgene, 80189). Cells were selected using 0.5μg/mL puromycin (Millipore Sigma, P9620). To activate expression of Cas9, cells were treated with 1μg/mL doxycycline (Millipore Sigma, D5207) daily and harvested after three days unless stated otherwise.

### Cell lysis

For immunoblots, whole cell extracts were generated by scraping then collecting cells via centrifugation. Cells were washed twice with ice-cold phosphate buffered saline (PBS) and lysed with RIPA buffer (Sigma), supplemented with Halt Phosphatase and Protease inhibitors (Pierce), 1mM phenylmethylsulfonyl fluoride (PMSF, Sigma), and 25U/mL Universal Nuclease (Pierce). Extracts were incubated on ice, and protein levels were quantified using BCA Assay (Pierce) and supplemented with 4x Laemmli Buffer with 2-mercaptoethanol (Biorad). Samples were boiled and sonicated. To generate histone extracts, cells were scraped then centrifuged and washed twice with ice-cold PBS. Cells were incubated with hypotonic lysis buffer (10mM Tris-HCl, pH 8.0, 1mM KCl, 1.5mM MgCl2, 1mM DTT, Halt, and 1mM PMSF) for 30 minutes on rotator at 4°C. Nuclei were pelleted at 10,000g for 10 minutes at 4°C, and histones were extracted from pelleted nuclei using 0.2N HCl at 4°C overnight on rocker. Nuclear debris was pelleted at 16,000g for 10 minutes, and histone extract was quantified using BCA Assay, neutralized with an equal volume of 1M Tris-HCl, pH8.0, and supplemented with 4x Laemmli Buffer then boiled. To fractionate chromatin, cells were resuspended in Buffer A (10mM HEPES (pH 8.0), 10mM KCl, 1.5mM MgCl2, 0.34M sucrose, 10% glycerol, 1mM DTT, 0.1% Triton-X100 and protease and phosphatase inhibitors described above) and incubated on ice for 10 minutes. Nuclei were pelleted at 1,300g for 5 minutes at 4°C. Supernatant was removed and clarified at 20,000g for 15 minutes at 4°C to remove insoluble debris. Nuclei were washed again in Buffer A, then lysed in Buffer B (3mM EDTA, 0.2mM EGTA, 1mM DTT and protease and phosphatase inhibitors). Insoluble chromatin was collected by centrifugation at 1,700g for 5 minutes at 4°C. Chromatin was washed in Buffer B, centrifuged then resuspended in Laemmli Buffer supplemented with Universal Nuclease, 2-mercaptoethanol and sonicated.

### Immunoblots

Protein extracts were run out using 4-20% Criterion TGX Precast Gels (Biorad) then transferred using Trans-blot Turbo Transfer System (Biorad) for semi-dry (histones and low-molecular weight proteins) or Criterion Blotter for wet transfer (high-molecular weight proteins). Proteins were transferred onto PVDF membranes (ThermoFisher), blocked with 5% BSA in TBS-Tween 0.1% (TBSTw) then probed for antibodies in block buffer. SuperSignal West Pico PLUS or Femto Maximum chemiluminescent solution (Pierce, 34580 and 34095) was added and blots were imaged using either film (HyBlot) or BioRad Gel Doc. Primary antibodies used were anti-SETD2 (Sigma, HPA042451), H3K36me3 (Cell Signaling, 4909), β-actin (Cell Signaling, 4970), Histone 3 (Cell Signaling, 14269), Cas9 (Active Motif, 61978), RAD51 (Millipore, 05-530-1), RAD52 (Santa Cruz Biotechnology, sc-365341), γH2A.X phosphor-Serine139 (Millipore, 05-636), and β-tubulin (Clone E7, gift from K. Verhey). Secondary antibodies used were anti-rabbit and anti-mouse HRP conjugates (Pierce, W401B and W402B).

### Cell and metaphase spread immunofluorescence and imaging

Metaphase spreads for immunofluorescence were prepared as described previously^42^. In brief, cells were cultured on cleaned glass coverslips (No. 1.5) in multi-well plates or glass-bottom multi-well plates (CellVis). Cells were fixed either by ice-cold 100% methanol for 10 minutes or 4% paraformaldehyde in Cytoskeleton Buffer (100mM NaCl, 300mM Sucrose, 10mM PIPES pH6.8, 3mM MgCl2, 0.5% TritonX100). Cells were rinsed twice with PBS, incubated with PBS with 0.1% TritonX100 (PBSTx) for 10 minutes. Cells were blocked with 2% BSA in PBSTx, then stained with primary and secondary antibodies in blocking buffer. Cells were washed with PBS, counterstained with Hoechst (2.5μg/mL) then mounted with ProLong Antifade Gold or stored in PBS.

For immunofluorescence of metaphase chromosomes, cells were treated with 150ng/mL of colcemid (KaryoMAX, ThermoFisher) for 1-2 hours then harvested via shake-off then pelleted via centrifugation. Cells were gently resuspended and swelled with 1:1 ratio of 75mM KCl:0.9% Sodium Citrate (MEFs) or 75mM KCl (HeLa), prewarmed to 37°C. Cells were incubated for 5-15 minutes then cytospun onto poly-lysine coated slides at 1300rpm for 10 minutes. Cells were extracted using KCM buffer (10mM Tris pH8.0, 120mM KCl, 20mM NaCl, 0.5mM EDTA, 0.1% TritonX100) for 10 minutes. Samples were blocked using 1% BSA in KCM, stained with antibodies in 1% BSA in KCM, then fixed with 4% PFA in KCM for 10 minutes. Chromosomes were rinsed with PBS then counterstained with DAPI (1μg/mL) in PBS then mounted with VectaShield (H100) and sealed. Primary antibodies used were anti-αTubulin (12G10, gift of K. Verhey), cGAS (Cell Signaling, 31659), RPA (Cell Signaling, 2208), CENP-A (Cell Signaling, 2048), CENP-A (Abcam, ab13939), HEC1 (Santa Cruz Biotechnology, sc-135934), HEC1 (GeneTex, GTX70268), H3K36me3 (Abcam, ab9050), and CENP-B (Santa Cruz, sc-32285). Secondary antibodies used were anti-mouse and anti-rabbit AlexaFluor conjugates (ThermoFisher, A21121, A11001, A21236, A11032, A32733, A11034, A11037, and A21135).

Images of mitotic cells and metaphase spreads were acquired on Deltavision Elite imaging system (GE Healthcare) equipped with a Cool SnapHQ2 charge-couple device (CCD) camera (Roper), using a 60x 1.4 numerical aperture (NA) or 100x 1.4 NA objectives (Olympus). Images were acquired using pre-set Alexa filter settings, and optical sections were collected at 200-nm intervals and processed using ratio deconvolution in softWoRx (GE Healthcare, version 6.5.2). Images were prepared for publication using FIJI (Version 2.1.0/1.53o). Images of mitotic and interphase cells are maximum intensity projects, and metaphase spreads are single z-slices. Images of interphase cells for quantifying nuclear phenotypes were acquired using Cytation 5 (Biotek) using 20x 0.45 NA objective (Olympus) with standard filter sets. Nuclear images and metaphase spreads were manually quantified.

### Fluorescence *in situ* hybridization

Cells were treated with KaryoMAX then harvested via trypsinization. Cells were centrifuged then gently resuspended in 1:1 ratio of 75mM KCl: 0.9% Sodium Citrate (MEFs) or 75mM KCl (HeLa), for 5-15 minutes. Cells were partially fixed by addition of 1/10 volume of ice-cold Carnoy’s fixative (3:1 methanol:acetic acid). Cells were centrifuged, and gently resuspended then fixative was added, dropwise while vortexing. Cells were incubated at least 30 minutes on ice then centrifuged and resuspended in ~0.5mL fixative then dropped onto ice-cold slides. Slides were air-dried at room temperature overnight then stored at 4°C. Slides were re-hydrated in 2X SSC for two minutes, dehydrated in ethanol series (2 minutes each in 75%, 90%, 100% EtOH) then air dried. Centromere (PNA Bio, CENPB-AF488 or −647), telomere (PNA, TelC-A647, 1:500), Acro-P (Cytocell, LPE NOR, undiluted) or rDNA (Empire Genomics, RPCI23-225M6, 1:5) FISH probes were diluted in hybridization solution (10mM Tris-HCl pH7.2, 70% formamide, 0.5% blocking reagent) then added to slide. Slides were denatured at 78°C for 2-5 minutes, then incubated for overnight at 37°C. Slides were washed in 0.4X SSC then 2X SSC with 0.5% Tween-20 for two minutes each. Slides were washed three times with PBS adding DAPI (1μg/mL) to second wash. Slides were dehydrated using ethanol series then mounted with ProLong Antifade Gold (Invitrogen). If spreads were subjected to FISH following immunofluorescence, the immunofluorescence protocol was used, and spreads were dehydrated using an ethanol series prior to FISH protocol. If spreads were stained only with DAPI, the denaturing and hybridization steps were omitted. For G-banding and Q-FISH, samples were prepared, imaged and analyzed by Creative Bioarray.

### Multicolor FISH

*Setd2^flox/flox^* MEFs were treated with vehicle or 3 μM 4-OHT for three days, and mitotic cells were enriched by incubation in 100 ng/mL colcemid (KaryoMAX) for ~5 hours. To prepare metaphase chromosome spreads, cells were collected by trypsinization, swollen with pre-warmed 0.075 M KCl at 37°C for 6 minutes, and fixed by washing and resuspending in ice-cold Carnoy fixative. Metaphase spreads were then dropped onto slides and allowed to air dry. For multicolor FISH, 3 uL of 21XMouse probes (MetaSystems) were applied to the sample and sealed with a coverslip using rubber cement. Samples and probes were co-denatured at 75°C for 2 min followed by incubation at 37°C overnight in a humidified chamber using ThermoBrite System (Leica). Slides were then washed for 2 min in 0.4x SSC at 75°C followed by a 30 second wash in 2x SSC at room temperature. Slides were counterstained with DAPI and mounted with an anti-fade solution. Metaphases were imaged using a Metafer Slide Scanning Platform (MetaSystems). Slides were first scanned to search for metaphase spreads using MSearch and then automatically captured using Autocapt at 63X magnification. Multi-color karyotypes were generated using Isis (v5.8.12, MetaSystems) and adjusted for imaging threshold. Dicentric chromosomes were identified as chromosomes harboring two DAPI-bright staining regions, and inter-chromosomal rearrangements were assessed by the appearance of different colored FISH probes hybridizing to the same dicentric chromosome.

## Figure Legends

**Extended data Figure 1:**
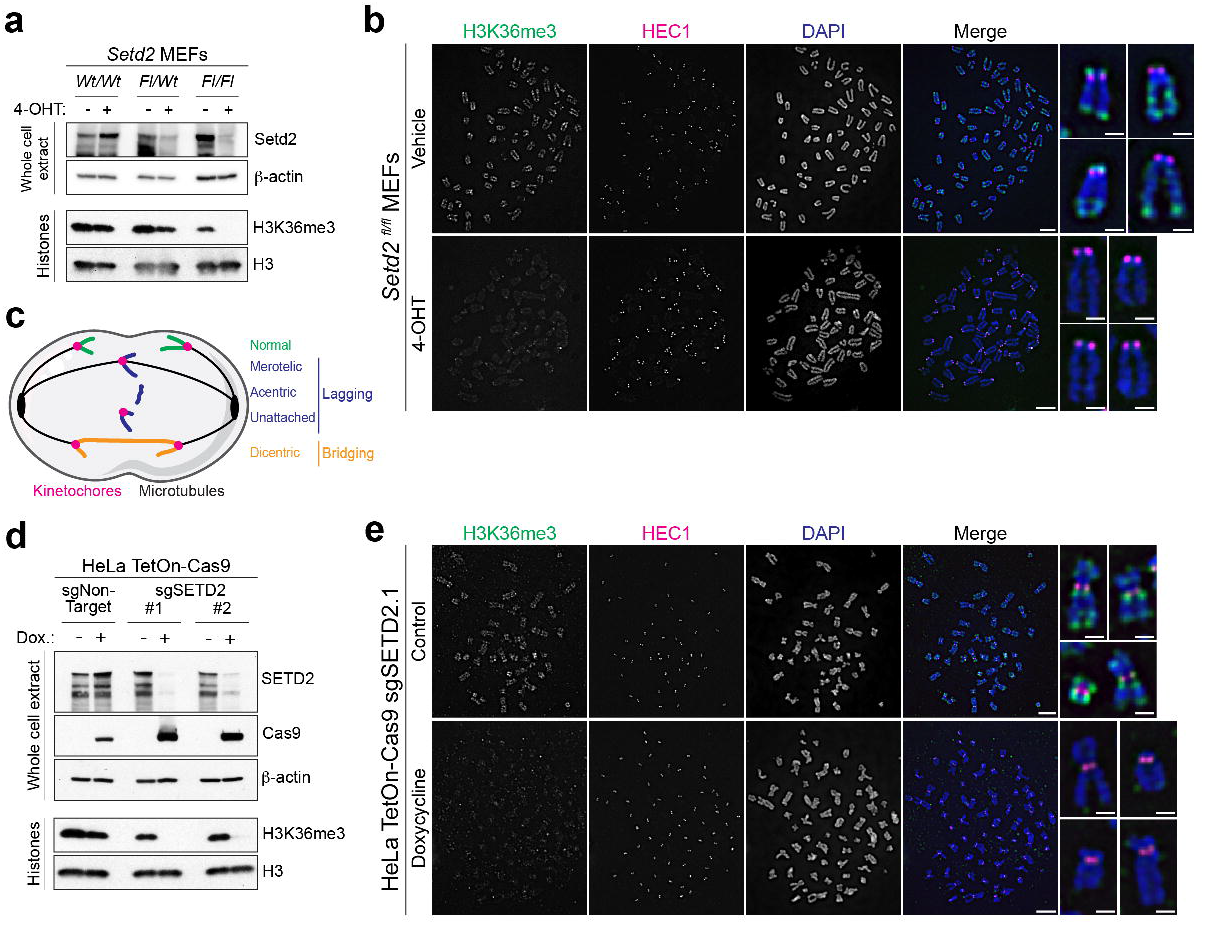
SETD2 is required for H3K36me3, which is present along chromosome arms, sub-telomeric and pericentric regions. **a,** Western blot of MEFs with no, one or two floxed alleles of *Setd2* (*Wild-type/Wt*, *Flox/Wt, Flox/Flox*, respectively). 4-OHT leads to deletion of *Setd2* allele(s) and a concurrent decrease in H3K36me3. Top panels are whole cell extract, acid-extracted histones at bottom. **b,** Metaphase spreads from *Setd2^fl/fl^* vehicle (EtOH) or 4-OHT treated MEFs, immunostained for H3K36me3, HEC1 (kinetochore protein) and DAPI (DNA). Scale bars are 5μm at left, and 1μm for individual chromosomes at right. **c,** Model of different types of chromosome mis-segregation. **d,** Western blot of HeLa TetOn-Cas9 cells expressing sgNon-targeting or sgSETD2. Doxycycline is added to activate expression of Cas9, leading to deletion of SETD2 and loss of H3K36me3. Top panels are whole cell extract, acid-extracted histones at bottom. **e,** Metaphase spreads from *SETD2* sgSETD2.1 HeLa cells, control or doxycycline treated, immunostained for H3K36me3, HEC1 (kinetochore protein) and DAPI (DNA). Scale bars are 5μm at left, and 1μm for individual chromosomes at right.

**Extended data Figure 2.**
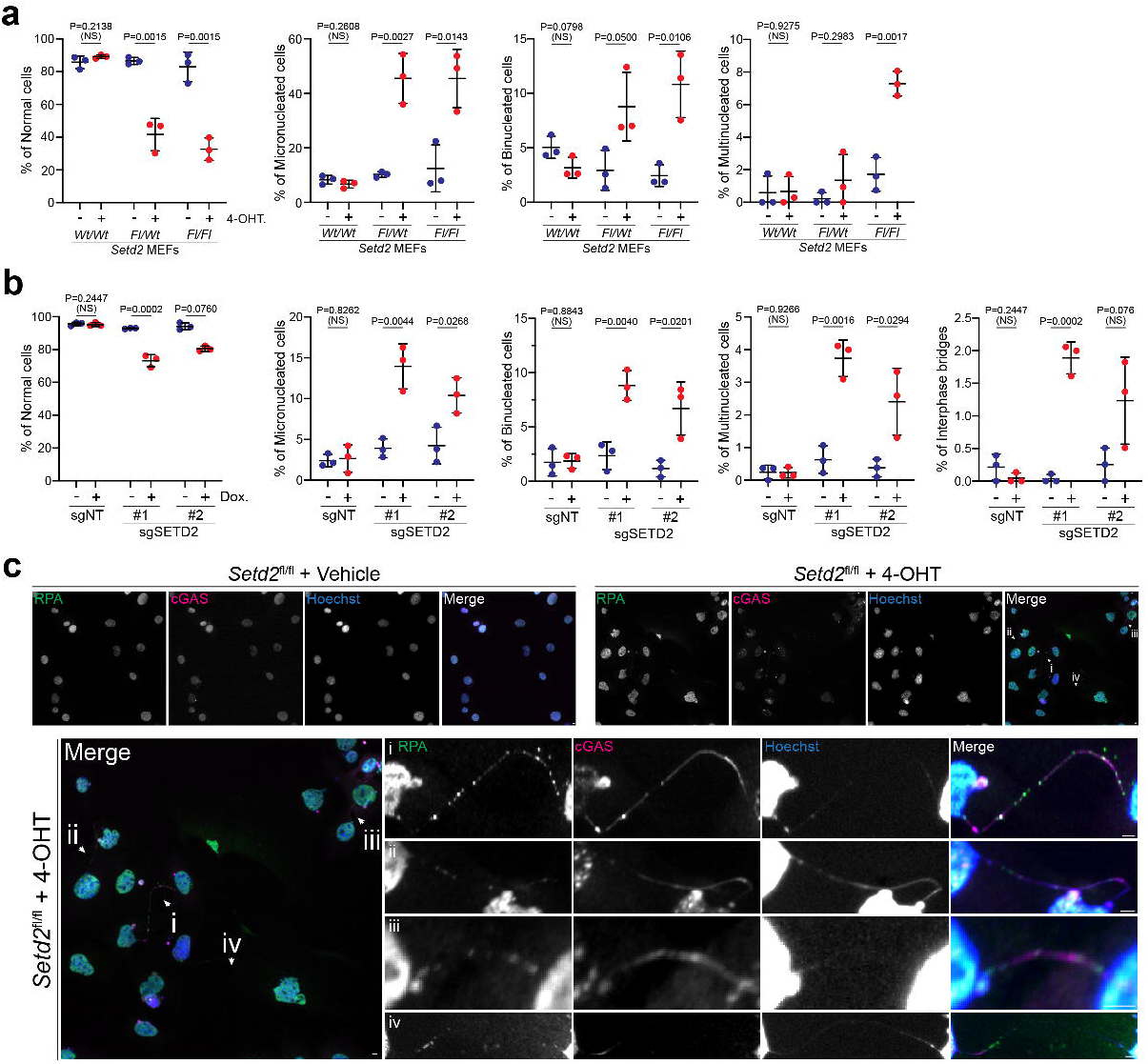
Setd2 loss promotes micronucleation, multinucleation, binucleation and bridging. **a,** Quantification of nuclear phenotypes of MEFs described in **a**, where *Setd2* deletion leads to less normal nuclei, and more micronucleated, binucleated, and multinucleated cells. N=3 independent experiments **b,** Quantification of nuclear phenotypes of HeLa cells described in **c**, where *SETD2* deletion leads to less normal nuclei, and more micro-, bi-, and multi-nucleated cells as well as an increase in cells with interphase bridges. N=3 independent experiments. For each plot, error bars are s.d. and P-values derived from unpaired t-test between each condition within a genotype. **c,** Image of *Setd2^flox/flox^* MEFs treated with vehicle (control, left) or 4-OHT (*Setd2* knockout, right) demonstrating that there are more interphase bridges in cells lacking *Setd2*. RPA (green) and cGAS (magenta) are markers of extranuclear DNA and bridges. Scale bars are 5μm.

**Extended data Figure 3:**
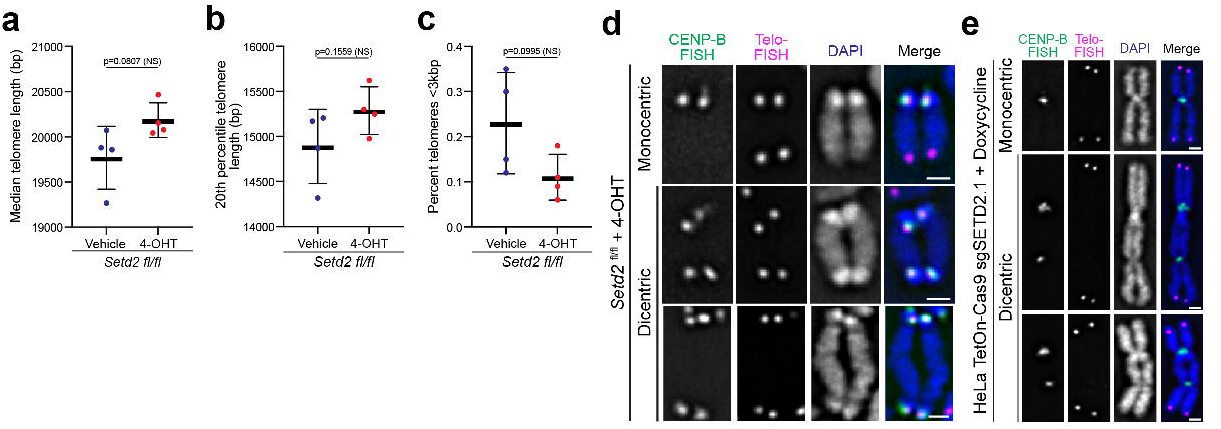
Telomere attrition or fusion does not occur following acute *Setd2* deletion. **a,** Quantification of median telomere length (base pairs, bp) in vehicle and 4-OHT-treated *Setd2^fl/fl^* MEFs. n=4 technical replicates. **b,** Quantification of length of 20^th^ percentile in vehicle and 4-OHT-treated *Setd2^fl/fl^* MEFs. n=4 technical replicates. **c,** Quantification of percent of telomeres less than 3 kilobase pairs long (kbp) in vehicle and 4-OHT-treated *Setd2^fl/fl^* MEFs. n=4 technical replicates. **d,** Representative images of CENP-B box and telomeric FISH on metaphase spreads from 4-OHT-treated *Setd2^fl/fl^* MEFs. Telo-FISH signal is not present between centromeres, suggesting lack of telomeric fusion. Scale bars are 1μm. **e,** Representative images of CENP-B box and telomeric FISH on metaphase spreads from doxycycline-treated HeLa TetOn-Cas9 sgSETD2.1 cells. Telo-FISH signal is not present between centromeres, suggesting lack of telomeric fusion. Scale bars are 1μm. Error bars are s.d. and P-values derived from unpaired t-tests.

**Extended data Figure 4:**
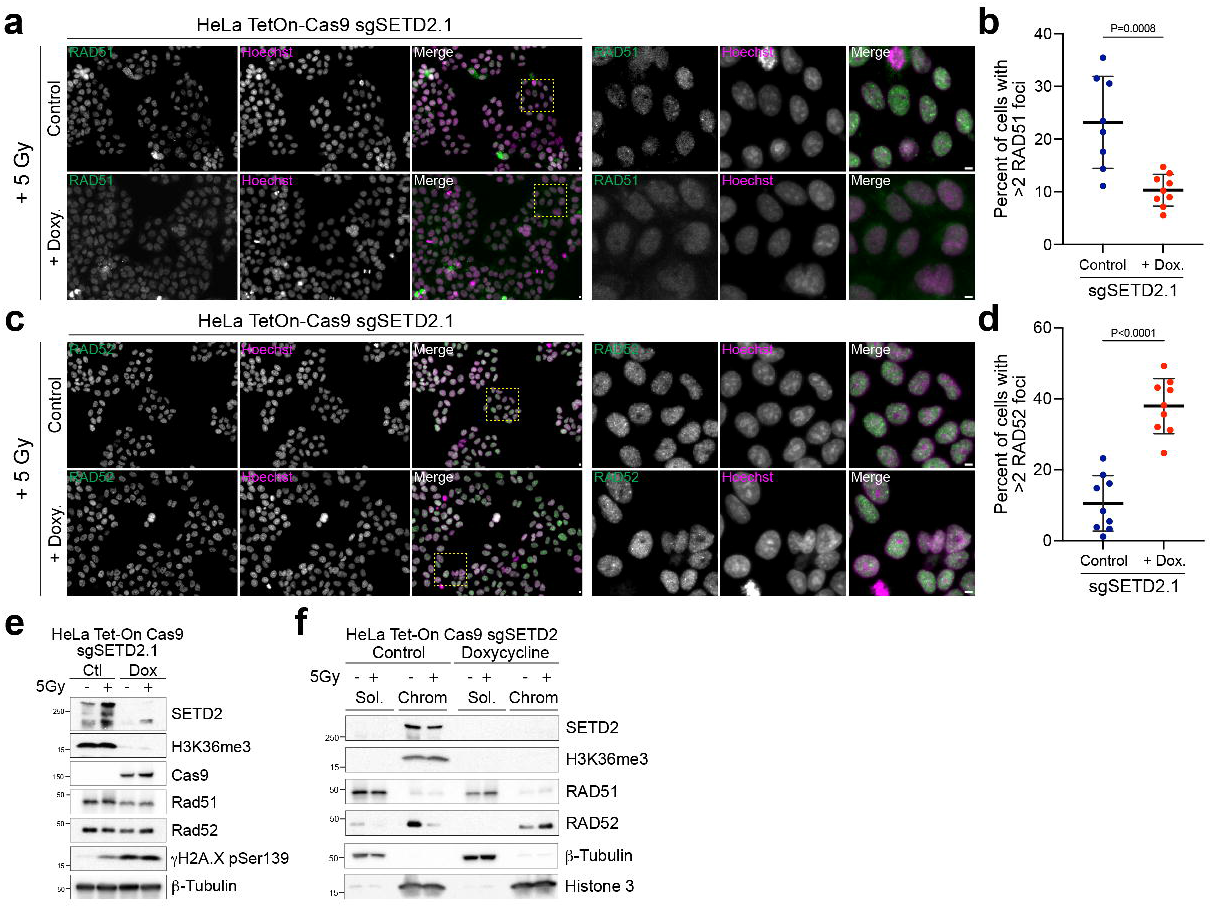
*SETD2*-knockout cells utilize RAD52 over RAD51 following DNA damage. **a,** Images of control or doxycycline-treated HeLa TetOn-Cas9 sgSETD2.1 cells were irradiated with 5Gy, recovered for 4 hours, then fixed and stained for RAD51 and Hoechst (DNA). Scale bars are 5μm. **b,** Quantification of percentage of cells with more than 2 RAD51 foci in control (n=8 technical replicates) or doxycycline-treated (n=9 technical replicates) HeLa TetOn-Cas9 sgSETD2.1 cells described in **a**. Data from one experiment and representative of two biological replicates. Error bars are s.d. and P-values derived from unpaired t-tests. **c,** Images of control or doxycycline-treated HeLa TetOn-Cas9 sgSETD2.1 cells were irradiated with 5Gy, recovered for 4 hours, then fixed and stained for RAD52 and Hoechst (DNA). Scale bars are 5μm. **d,** Quantification of percentage of cells with more than 2 RAD52 foci in control (n=9 technical replicates) or doxycycline-treated (n=9 technical replicates) HeLa TetOn-Cas9 sgSETD2.1 cells described in **c**. Data from one experiment and representative of two biological replications. Error bars are s.d. and P-values derived from unpaired t-tests. **e,** Immunoblots of whole cell extracts from control or 5 Gy irradiated cells described in **a**. While SETD2 and H3K36me3 are lost and a slight increase in DNA damage (γH2AX phosphor-Ser139) occurs in doxycyline treated cells, other protein levels remain unchanged. **f,** Immunoblots of soluble and chromatin-bound fractions from cells described in **a**. While chromatin loading of RAD52 is normally decreased following irradiation (5 Gy), *SETD2*-knockout cells have more chromatin-associated RAD52.

## Acknowledgements

We thank Brian Strahl, Jared Nordman, Jeffrey Rathmell and members of the WKR laboratory for their constructive input; I. Cheeseman (MIT), K. Verhey (U. of Michigan), and J. Rathmell (VUMC) laboratories for reagent and resource sharing; the Cell and Developmental Biology Equipment Resource (Vanderbilt University), Cell Imaging Shared Resource (VU) and Vanderbilt Ingram Cancer Center for imaging and technical support (P30CA068485). This work was supported by NIH 5T32CA009592 (L.V.), US DOD W81XWH221076 (P.L.), DOD KC200259 (F.M.M.), KCA W81XWH-21-1-0786 (F.M.M.), NIH R01CA20301 (W.K.R., C.L.W.), NIH R01CA275082 (W.K.R., R.D.), and CPRIT RP220332 (R.D.). Extended data Fig. 1c, Fig. 4j were modified from “Mitosis” by Servier Medical Art, licensed under a Creative Commons Attribution 3.0 unported license. Fig. 3d is modified from illustrations on Bioicons.

## Contributions

F.M.M. and W.K.R. conceived this project, with input from R.O., C.A.L., Ru.De., and C.L.W. F.M.M., E.S.K., A.T., Ra.Da., L.V., and S.R.N. designed, performed and analyzed the experiments. T.G. performed cytogenetic analyses of Geimsa banding (Fig. 4c). F.M.M. and W.K.R. acquired funding and supervised this project. F.M.M. and W.K.R. wrote the manuscript, with input from all authors.

## Authors Addresses

Frank M. Mason, Emily S. Kounlavong, Anteneh Tebeje, Logan Vlach, & W. Kimryn Rathmell: 1161 S 21^st^ St. MCN Suite D3100, Vanderbilt University Medical Center, Nashville, TN 37232.

Rashmi Dahiya & Peter Ly: UT Southwestern Medical Center, 5323 Harry Hines Blvd., Dallas, TX 75390.

Stephen Norris: 1170 Veterans Blvd., South San Francisco, CA 94080.

Tiffany Guess: Molecular Pathology Laboratory Network, Inc. 250 E. Broadway, Maryville, TN 37804. Courtney A. Lovejoy: 2215 Garland Ave., Vanderbilt University, 613 Light Hall, Nashville, TN 37232. Ruhee Dere: Center for Precision Environmental Health, One Baylor Plaza Alkek Tower N1317.04, Houston, TX 77030.

Cheryl L. Walker: Center for Precision Environmental Health, One Baylor Plaza Alkek Tower N1317.01, Houston, TX 77030.

Ryoma Ohi: 3065 BSRB, 109 Zina Pitcher Place, Ann Arbor, MI 48109.

